# Optimized Detection and Inference of Names in scRNA-seq data

**DOI:** 10.1101/2025.01.27.634698

**Authors:** Janyerkye Tulyeu, David Priest, James B. Wing, Jonas Nørskov Søndergaard

## Abstract

Accurate identification of immune cell subsets in single-cell (sc)RNA-seq data is critical for understanding immune responses in autoimmune diseases, infections, and cancer. One caveat of scRNA-seq is the inability to properly assign rare immune cell subsets due to gene dropout events. To circumvent this caveat, we here developed Optimized Detection and Inference of Names in scRNA-seq data (scODIN). scODIN uses an informed holistic two-step approach combining expert knowledge with machine learning to rapidly assign cell type identities to large scRNA-seq dataset. First, scODIN uses key lineage-defining markers to identify a set of core cell types. Second, scODIN compensates for dropout events by integrating a k-nearest neighbors (kNN) algorithm. We additionally programmed scODIN to detect dual and transitional phenotypes, which are usually overlooked in conventional analyses. Consequently, scODIN may enhance our understanding of immune cell heterogeneity and provides comprehensive insights into immune regulation, with broad implications for immunology and personalized medicine.

## Introduction

Accurate identification of immune cell subsets is important for understanding their roles in various immune processes, such as autoimmunity, infections, and cancer. Traditionally, methods such as flow cytometry and mass cytometry have been used to characterize immune cell populations based on specific marker expression^1,2^. While these techniques have provided valuable insights, they are limited by the number of markers that can be analyzed simultaneously. The advent of single-cell RNA sequencing (scRNA-seq) has revolutionized the ability to profile gene expression at the single-cell level, offering unprecedented insights into the heterogeneity of immune cell populations^3,4^. This technology has allowed researchers to uncover previously unrecognized cell states and subsets, leading to a more nuanced understanding of immune responses^3–6^. Despite this, it remains valuable to place scRNA-seq data into conceptual boxes such as T helper (Th)-cell subsets (e.g. Th1, Th2), to provide a structure in which deeper insights can be validated. Unfortunately, scRNA-seq poses significant challenges for accurate cell subset identification, primarily due to the phenomenon of single-cell “dropout events”, where genes, especially those with low expression levels, are not detected in all cells^7,8^. This issue, together with transcriptional bursting ^9^, complicates the identification of key markers, such as transcription factors, chemokine receptors, and cytokines, that is used to define T-helper and T regulatory (Treg) subsets^10–14^. Recent efforts to enhance cell type determination by integrating protein marker expression with gene expression data have shown promise^15,16^. However, these methods still struggle to accurately assign T cells to their correct subsets^17^. To address these limitations, we here developed Optimized Detection and Inference of Names in scRNA-seq data (scODIN), a novel computational approach designed to improve the accuracy and speed of immune cell subset identification. scODIN builds on existing algorithms^18–21^ by implementing a tiered system that prioritizes the identification of rare cell types before assigning broader, more generic labels. Additionally, scODIN incorporates a k-nearest neighbors (kNN) algorithm^22,23^ to extend the identification of cells not clearly identified in the first step due to dropouts or unclear expression of key markers. kNN identifies the nearest neighbors of each cell based on their gene expression profiles and assigns cell types based on the similarity to these neighbors^23^.

One of the most challenging aspects of T cell subset identification is the detection of mixed phenotypes, such as Th1/Th17 cells^24^. These cells co-express markers of both Th1 and Th17 subsets, making them difficult to classify. However, mixed phenotypes may be important in understanding immune responses in various diseases, including autoimmune disorders and infections^25–28^. scODIN addresses this challenge by allowing double labels of cells with known or unknown mixed phenotypes. Another significant advancement offered by scODIN is the identification of intermediate phenotypes. These are cells that exhibit characteristics of two distinct subsets, such as those that exist between naïve and central memory T cells^29,30^. Intermediate phenotypes are often overlooked in conventional analyses because they do not fit neatly into predefined categories. However, they may provide a more dynamic picture of our immune response and may play critical roles in immune regulation and disease progression, making their accurate identification essential for a complete understanding of immune responses. By identifying these intermediate states, scODIN provides a more comprehensive view of T cell heterogeneity and offers insights into the dynamic nature of immune cell differentiation. Finally, in order to make scODIN as user-friendly as possible, we have coded it to work as an add-on to the R package Seurat, a widely used toolkit for scRNA-seq data analysis^31^.

In this study, we present the development and validation of scODIN, demonstrating its superior performance compared to existing algorithms in terms of both speed and accuracy. We also highlight its ability to accurately identify rare T cell subsets and mixed/intermediate phenotypes, which are critical features for applications in immunology and personalized medicine. While the example framework given is for CD4 T-cells, these methods can also be applied to all other cell types by adapting the same approach.

## Results

### scODIN: Optimized Detection and Inference of Names in scRNA-seq data overview

There are currently two main approaches to cell type classification in single-cell RNA-seq data. Fully automated or expert-input manual classification. scODIN uses the best of both approaches and enables fast and accurate classification of immune cells in single-cell RNA-seq data (**Fig. 1**). Usually, the top level lineage classification in PBMCs or similar tissue is straightforward for the expert eye to classify, as the diversity of cell types is high. Nevertheless, no highly accurate approach has been developed to automate this process yet, and expert annotation is still time-consuming, thus calling for a fast, but most importantly, accurate approach. Therefore, we developed scODIN to conduct this first step automatically (**Fig. 1a, Fig. S1**). As input, scODIN requires the normalized and scaled scRNA-seq dataset, a gene priority table, and an accepted double labels table (**Fig. S1a**). Further explanation of how the scODIN score is calculated can be seen in **Fig. S1b**. The second layer classification of rare immune subsets is a far from trivial task and hard to achieve in a single step. Thus, we programmed scODIN to conduct this difficult task using a 2-step approach (**Fig. 1b, d**) with an optional 3rd step allowing additional detailed phenotyping of double labelled cells such as Th1/Th17 cells, and intermediate phenotype cells, such as ones in-between naïve and central memory cells (**Fig. 1c**). To enable thorough and accurate determination of rare cells, we developed a tier-based system to first identify rare cells (such as Tfr, identified by CXCR5+Foxp3+) in tier 1 and subsequently common and more generic categories of cell types such as central memory (CM) in tier 2 (**Fig. 1b**). Finally, the identified cell types are expanded using kNN machine learning (**Fig. 1d**). In contrast to most other algorithms, scODIN runs very fast on large single-cell RNA-seq dataset. For instance, starting from 647k cells the entire process takes around 6 minutes (**Fig. 1e**). In the next figures we will demonstrate how each of these steps works and which variables the user can adjust to fit their own dataset.

**Figure 1:**
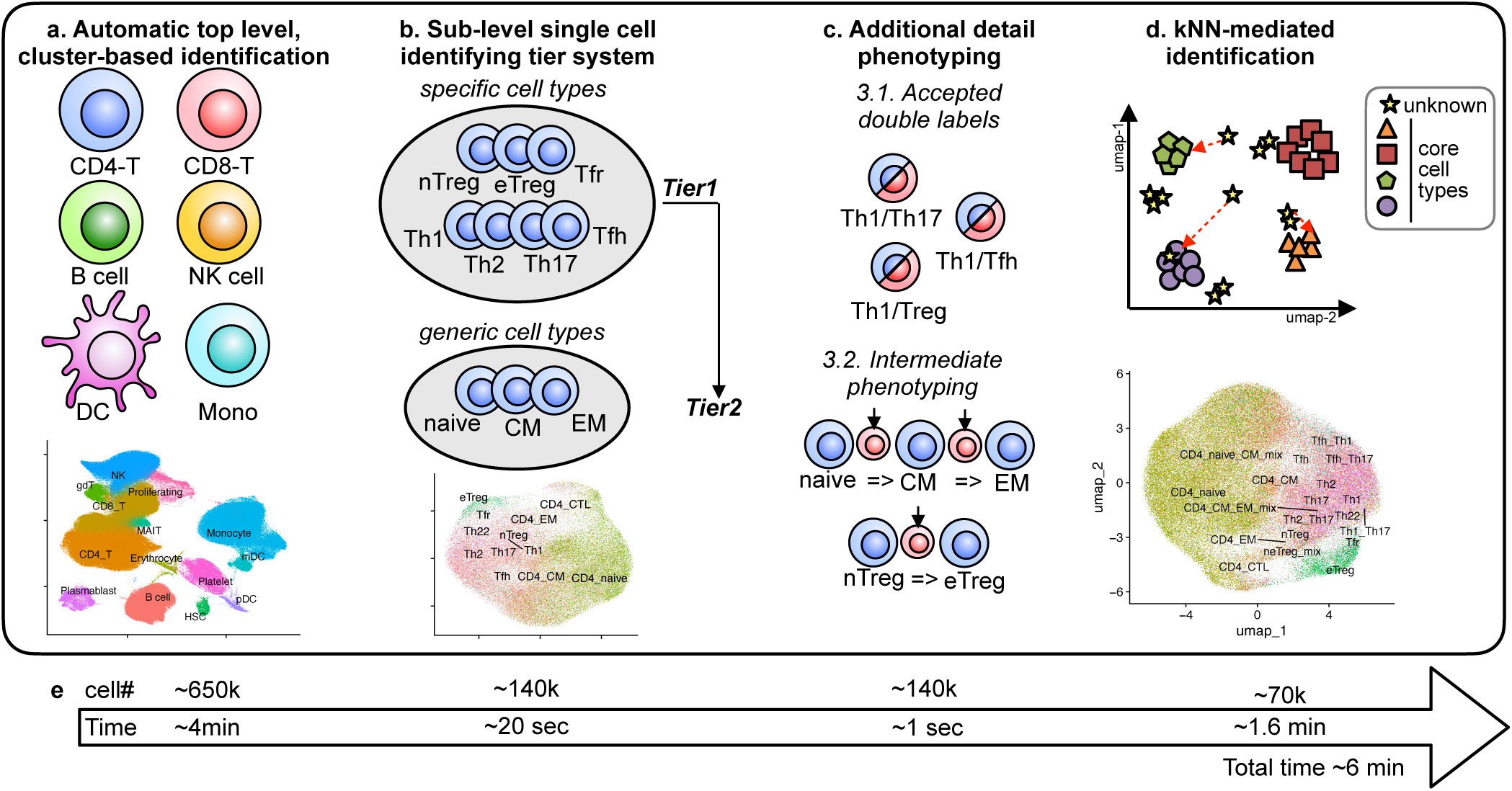
Optimized Detection and Inference of Names in scRNA-seq data (scODIN) overview. scODIN consists of three main steps and additional detail phenotyping. Above is a schematic of each step, in the middle, the UMAPs of identified cell types for each step, and below is an example of time used for the indicated cell numbers. **a**, Step one is a fast and accurate automatic cell type identification at the top level, to identify clusters of e.g. CD4 T cells (CD4-T), B cells, and monocytes (Mono) for further subset phenotyping. At the top level, the cell type identities are made on a cluster level. The remaining steps in scODIN is truly single-cell determination of cell identities. **b**, A user-defined tier-based system is implemented to identify cells of different detail levels. In the example, specific subsets such as Tregs and T helper cells are identified in the first tier and more generic cell types such as central memory (CM) and effector memory (EM) are identified in the second tier. **c**, In the same calculation as b) it is possible to add accepted double labels, such as mixed T helper phenotypes or intermediate phenotyping, to identify cells that have an in-between gene expression of naïve and effector cells. **d**, Finally, the last step uses k-nearest neighbor (kNN) machine learning to expand cell identification in case crucial core genes have undergone single-cell dropout events. A user-defined cutoff of stringency is used to either give every single cell a category or leave some unidentified. **e**, Cell numbers and running times for each step of scODIN. Data in the example is from Stephenson *et al.*^32^

### scODIN top level automated classification outperforms other leading algorithms in terms of speed and accuracy

The first step in scODIN is a task that has been attempted by many other groups^18–21^. We thus benchmarked scODIN’s performance compared to these other algorithms (**Fig. 2**). Using datasets of varying size (3k, **Fig. 2a-f** to 650k cells, **Fig. 2g-j**) scODIN gave the same classification as the original authors gave with manual annotation. However, this should be understood in the context that, due to the 3-step approach in scODIN, there are no sub-classifications at the top level, making initial classification a simpler task. scODIN for instance will initially classify CD4 Tregs as CD4 T cells, and only at the next steps will the Tregs be identified. In contrast, all the other algorithms tested performed poorly in terms of accuracy. For instance, one algorithm classified CD8 T cells as CD8+ NKT-like cells, and all B cells as naïve B cells (**Fig. 2c**). In both cases these are the result of applying complex labels, representing subpopulations to the top-level data. Although the terms are similar by name it is biologically implausible that all CD8 T-cells would be NKT like or that all B-cells would be naive and should be accounted for as a misclassification. A few of the algorithms furthermore failed to run when a large single-cell RNA-seq dataset was used (with >100GB free RAM) (**Fig. 2k**). Thus, scODIN outperformed all other tested algorithms in terms of accuracy (**Fig. 2k**) and was close to 100% accurate on the tested dataset. In terms of speed, scODIN also outperformed all other algorithms, and could run a 647k cell dataset in 4 min, and a 3k dataset in less than one second (**Fig. 2e**).

**Figure 2:**
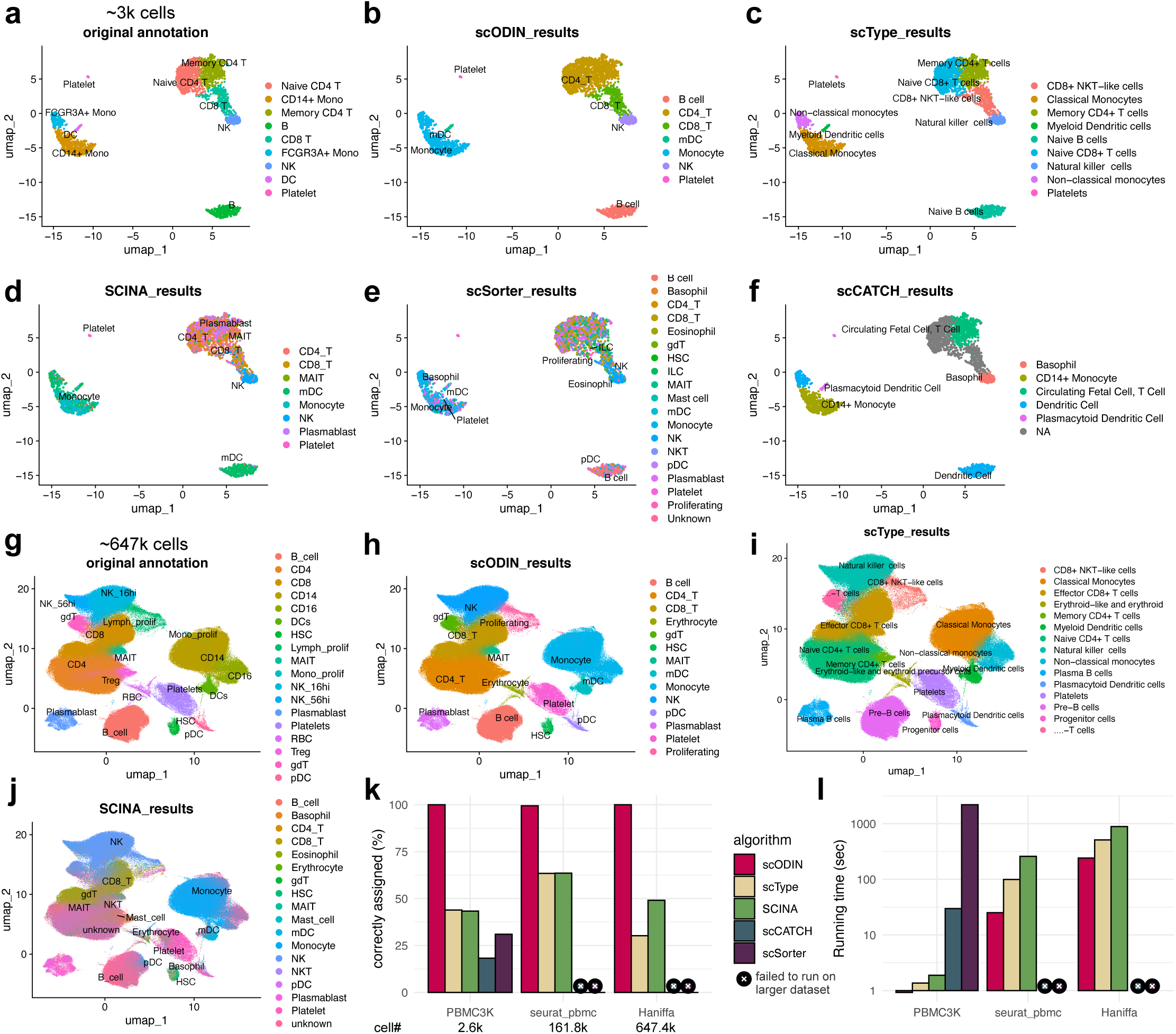
scODIN outperforms other leading algorithms in terms of speed and accuracy. **a-j**, UMAPs of automated cell type identification algorithms, using a ∼3k PBMC cell dataset^3^ (a-f), and a ∼647k PBMC cell dataset^32^ (g-j). **k-l**, Comparison of correctly assigned (given by the original expert annotated cell types) and running times between scODIN and other leading algorithms.

### scODIN tier-system enables thorough and accurate determination of rare cells

Using the 647k dataset subsetted to only CD4 T cells (∼140k cells) we next used the scODIN tier system to detect rare cell types (**Fig. 3**). In **Fig. 3a** and **Fig. S2a** we compare the number of cells detected for the classifications that overlapped with the original study by Stephenson *et al*.^32^. In the original study, Tregs were not sub-classified but using scODIN we could sub-classify these into nTreg, eTreg, and Tfr cells, and overall detected many more Tregs than the original study (**Fig. 3b-d, Fig. S2b**). Phenotypically the scODIN-identified Tregs all expressed FOXP3 (**Fig. 3f, g**) and some CTLA4, Helios (IKZF2), and CD25 (IL2RA) (**Fig. S2c-e**) confirming that this was not inflation by incorrect identification. In contrast, in the original study, Tregs were defined as all expressing FOXP3 and Helios, although 10-30% of Tregs do not express Helios^33^. The remainder of the FOXP3 expression in the original study was found mostly in naïve, CM, and Th22 cells (**Fig. 3b**). The scODIN tier-system made little difference for the detection of Tregs, however, scODIN with the tier-system detected more Tfh, Th1, Th2, and Th17 cells than scODIN without the tier system (**Fig. 3a**). For Tfh and Th22, scODIN detected less cells than the original study, however these cell types were most likely over-estimated in the original study, as key markers CXCR5 and IL22 respectively were missing in most of the cells (**Fig. 3b, h, k**). scODIN without the tier system performed better than the original study in terms of detected cell numbers and marker specificity (**Fig. 3c, f, i, l**). However, using scODIN with the tier system further improved the detection with more unknown cells expressing key markers being classified into specific cell types (**Fig. 3d, g, j, m**). This can clearly be seen by comparing CXCR5 expression without and with the tier system (**Fig. 3i** versus **3j**) or IL22 (**Fig. 3l** vs. **3m**). The reason the tier system works is because every cell gets a score for each cell type, and to classify a cell there is a competition of highest score (**Fig. S1c**). By removing irrelevant competition more cells can be classified.

**Fig. 3:**
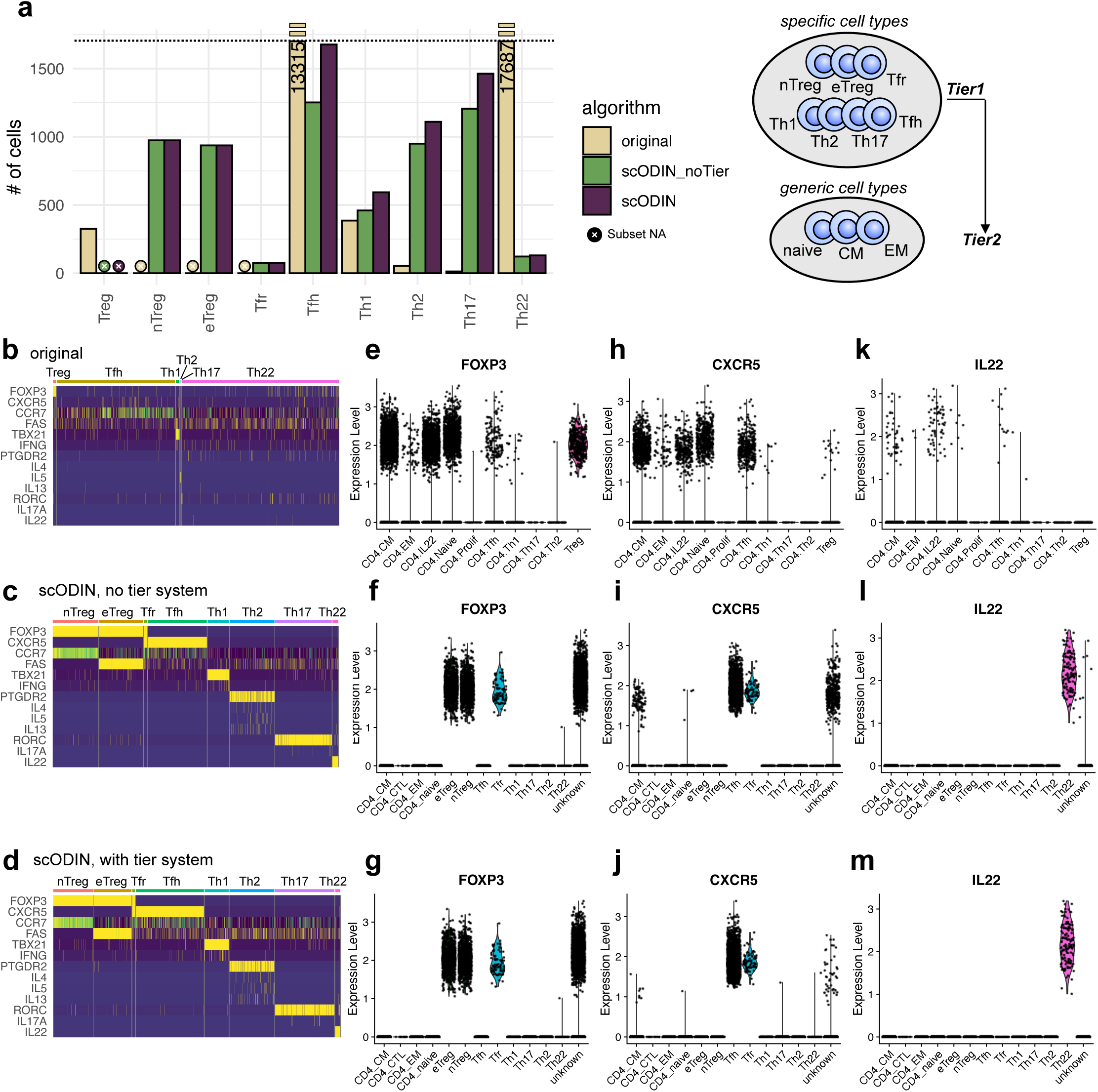
scODIN tier system enables thorough detection of rare cell types. **a**, Number of rare cell types detected by scODIN +/- the tier system compared to the original annotation^32^. The scale has been cut off at 1750, and bars with values above that have the number indicated on them. **b-d**, Heatmaps of key markers for rare immune cell subsets for b) the original annotation, c) scODIN without tiers, and d) scODIN with tiers. **e-m**, Violin plots of FOXP3 expression (e-g), CXCR5 expression (h-j), and IL22 expression (k-m) for the original annotation (first), scODIN without tiers (second), and scODIN with tiers (last).

### scODIN enables additional detailed phenotyping of mixed or intermediate phenotypes

Due to the nature of the scODIN scoring system, it is possible to retrieve double phenotype cells if the scores are within a certain threshold set by the user (**Fig. 4a-b**). For instance, if one cell has a similar score for Th1 and Th17, they will be labelled as Th1/Th17 cells (**Fig. S1c**). Using a similarity cutoff, the user decides the cutoff for single or double labels. A similarity cutoff of zero would yield no double labelled cells, while increasing similarity scores would yield more double labelled cells (**Fig. 4b**). As expected, double labelled Th1/Th17 and Th1/Tfh cells expressed TBX21 (T-bet) (**Fig. 4a**), and Th1/Th17, Th17/Th2, and Tfh/Th17 cells expressed RORC. Th17/Th2 cells additionally expressed PTGDR2 (CRTH2), and Th1/Tfh and Tfh/Th17 cells expressed CXCR5 (**Fig. 4a**). We also identified Tfr/nTregs and Tfr/eTreg double labelled cells (**Fig. 4a**), which we simplified to be labelled as Tfr using an optional “simplify_double_labels()” function in scODIN. In a similar principle, scODIN also allows identifying intermediate phenotypes, such as cells transitioning between naïve and CM (**Fig. 4c-e**) or between nTregs and eTregs (**Fig. 4f-h**). For CD4 T cells, we found cells starting from naïve going to naïve/CM through CM and to CM-EM had decreasing amounts of CCR7, SELL (CD62L), LEF1, and TCF7 and increasing amounts of CD69 and S100A4^34–37^ (**Fig. 4c**). Furthermore, FAS was expressed by CM and EM cells, while KLRG1 was mainly present at the EM stages^38^. Cells starting from naïve going to naïve/CM through CM and to CM-EM were furthermore moving from bottom to top on the UMAP (**Fig. 4d**) and having increasing pseudo time (**Fig. 4e**). However, pseudo time or UMAP localization did not distinguish CM-EM cells from EM. To further demonstrate the intermediate phenotype utility of scODIN, we also identified Treg cells transitioning between naïve and effector Tregs (**Fig. 4f-h**). All subsets expressed Foxp3 but going from nTreg through nTreg-eTreg to eTreg had decreasing CCR7 and TCF7 expression and increasing FAS expression (**Fig. 4f**). The density of cells moved from bottom to top of the UMAP (**Fig. 4g**), and pseudo time increased steadily through these scODIN-identified Treg subsets (**Fig. 4h**). In summary, scODIN enables identification of mixed and intermediate CD4 T cell phenotypes, which may allow a more detailed and dynamic picture of the immune response.

**Fig. 4:**
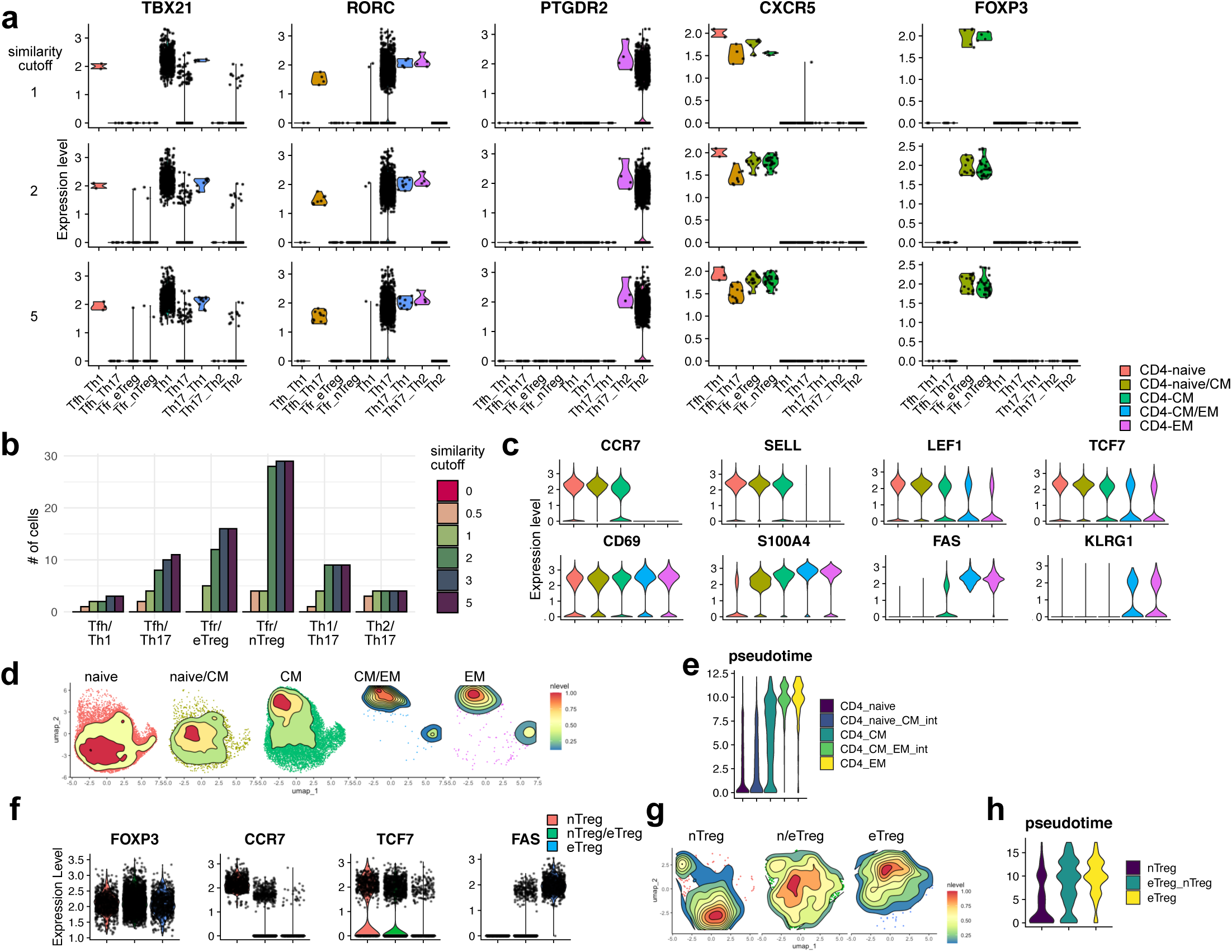
scODIN enables additional detailed phenotyping of mixed or intermediate phenotypes. **a-b**, scODIN can provide double labels if the cell has a mixed phenotype, such as simultaneous expression of TBX21 (Th1 marker) and RORC (Th17 marker). The user supplies a list of accepted double labels, and by adjusting a similarity cutoff more or less double labels are retrieved. **a**, Violin plots demonstrating key T cell subset markers (TBX21: Th1, RORC: Th17, PTGDR2: Th2, CXCR5: Tfh, FOXP3: Treg) on single and double-labeled cell types at different similarity cutoffs. **b**, Quantification of the number of double labelled cells at different similarity cutoffs. **c-e**, scODIN can also detect intermediate phenotypes, such as a cell transitioning between naïve and central memory or between central memory and effector memory. **c**, Violin plots demonstrating key distinguishing markers between naïve, central memory, and effector memory CD4 T cells on single and double-labeled cell types. **d**, UMAP showing different distributions of single and double-labelled cells. **e**, Violin plots showing pseudo time of each subset. **f-h**, Violin plots and UMAP demonstrating a transitional phenotype between nTreg and eTreg.

### Expansion of scODIN-identified core cells through kNN

In order to consider single-cell dropout events, we next used kNN to expand the number of identified cells, by using all scODIN core identified cells as the reference and all unidentified cells as the query dataset (**Fig. 5a-c**). As the core identified cells contained few rare cells, such as Tfr and Th1, but many CD4-CM and naïve cells, we first down-sampled the number of cells in each subset to have a comparable reference (**Fig. 5a**). As we in the previous steps identified some very rare cell types (<10 cells) it was undesirable to down-sample to the smallest subset. Thus, we instead down-sampled to the median-sized subset (937 cells). Next, all cells that did not receive a label in the core cell determination were used as the query dataset for kNN (**Fig. 5b**). These cells either had a low scODIN score, triple labels, or unaccepted double labels. In the standard kNN approach implemented by Seurat^22,23^ all cells receive an identification based on the highest score alone. As cells may have very similar or low scores, we here implemented a user-defined score filter, which will determine how many cells remain unassigned after kNN (**Fig. 5c**). If the user wants all cells to have an identity, the score filter can be set to 0 or it can be gradually increased to 1.0 leading to increasing number of unassigned cells (**Fig. 5c**). Which cutoff is proper can be judged based on expression of key markers (**Fig. 5d**). Since the initial step uses key genes such as FOXP3 this step will widen the definition to cells with Treg-like characteristics but relatively low levels of FOXP3 to account for dropouts and transcriptional bursting. While predicted Tregs did not express much FOXP3, they expressed other Treg markers such as CTLA4, IL2RA (CD25) and IKZF2 (Helios) (**Fig. 5d**). Similarly, while predicted Th1 did not express much TBX21 (T-bet), they expressed other Th1 markers such as IFNG, CXCR3 and STAT4 (**Fig. 5d**). kNN uses machine learning and dimensionality reduction to compare which unknown cells are most similar to core cells that have previously been identified. As the results are based on small differences in many genes simultaneously it is not possible to extract which other genes contributed to the identification decision. Instead, by comparing the expression of markers in the core identified cells (**Fig. 5e**) to the final labels generated by combining the core cells with the kNN-identified cells (**Fig. 5f**) it can be noted that key markers are still present after kNN although diluted compared to the core cell types. As the user may be interested in comparing results with and without kNN we have implemented information on the source of identification in the scODIN code. In summary, scODIN enables the expansion of cell type identifications to account for single-cell dropout events by virtue of kNN.

**Fig. 5:**
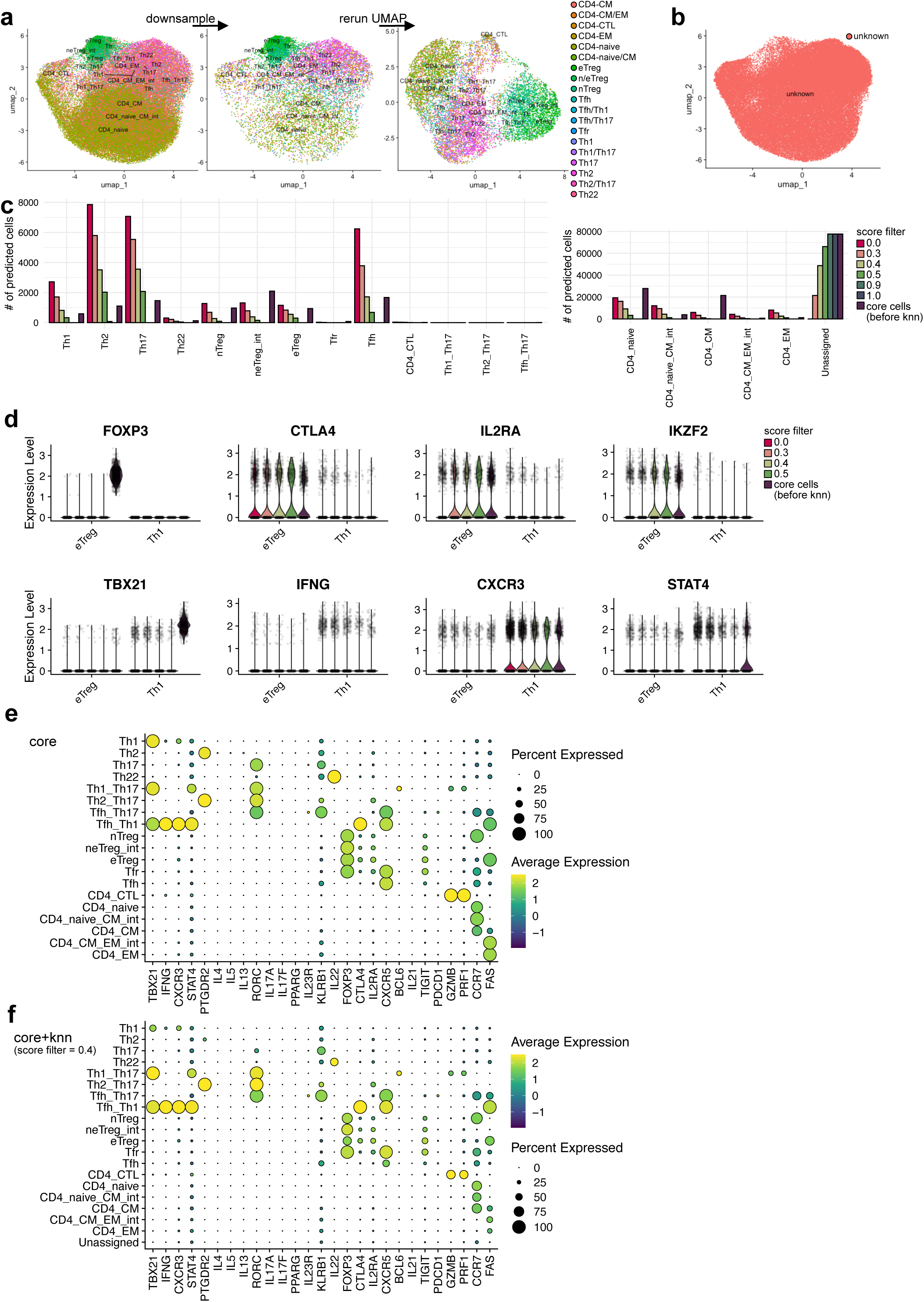
Extension of cell type identification by kNN. **a**, UMAPs of identified core cell types done in the previous steps of scODIN. Before kNN, the core cell types are down-sampled so that each cell type are represented by a similar number of cells. After down-sampling, the dimensionality reduction is rerun to generate a new UMAP. **b**, UMAP showing the cells that could not be identified in the previous steps of scODIN. These are the cells that will have identities predicted during kNN. **c**, Barplots showing the number of predicted cell types after kNN with different score filter cutoffs. For comparison, the number of cells identified as core cells in the first steps of scODIN are also displayed (purple bars). The graph is split in two due to varying numbers of cells for each type. **d**, Violin plots of Treg markers (FOXP3, CTLA4, IL2RA, IKZF2) and Th1 markers (TBX21, IFNG, CXCR3, STAT4) for cell types identified to be Treg or Th1 by scODIN’s kNN analysis. **e-f**, Dotplots of key marker expression in cells identified by the scODIN scoring system alone (e) and final labels combining core and kNN-identified labels (f).

### Through scODIN-mediated cell type identification, new biology may be discovered

Finally, to demonstrate the usability of the scODIN results, we correlated the identified cell types with COVID-19 severity (**Fig. 6, Fig. S3**). Using the updated cell identifications, we found a negative correlation with the frequency of Th1 with COVID-19 severity, as previously reported by others^39^ (**Fig. 6a-d**). Although the annotations done by Stephenson *et al.*^32^ also found a negative correlation, this was non-significant. Using scODIN annotations, the correlation of severity with Th22 was opposite to what could be found with the original annotation (**Fig. 6e-h**). Additionally, using scODIN we could gain information on intermediate phenotypes (**Fig. 6i-p**). The “standard” categories of CD4 naïve, CM and EM corresponded between the original study annotation and the scODIN-generated one. However, using scODIN we found that the intermediate phenotype of CD4-CM/EM cells were significantly different between healthy controls and COVID-19 patients (**Fig. 6o**). Similarly, the amount of total Treg gave similar results between original and scODIN annotations (**Fig. 6q-r**) but using the detailed subsetting information obtained by scODIN we found nTregs being significantly lower in COVID-19, and eTregs trending towards higher (p = 0.13) (**Fig. 6s-v**). Finally, the rare subset of CD4 CTL was found to be higher in COVID-19 using the scODIN annotation (**Fig. 6w**) as previously descibed^40,41^. Although results like that require further validation, scODIN provides the groundwork for new hypothesis generation in a fast and accurate manner.

**Fig. 6:**
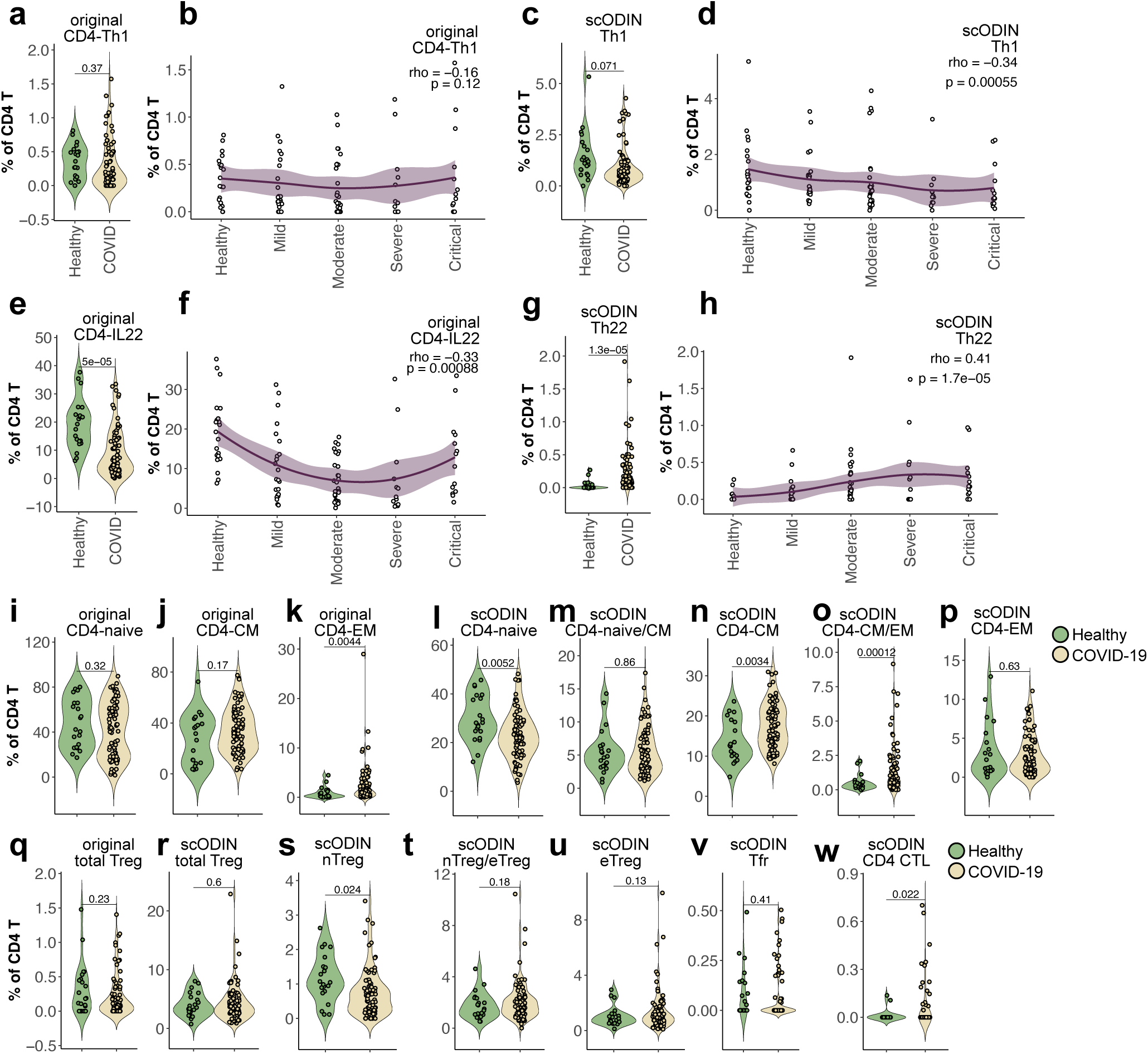
scODIN final annotations’ correlation with disease. **a**, Violin plot comparing healthy controls to all COVID-19 patients for originally identified Th1 cells by Stephenson *et al*.^32^ **b**, Correlation analysis of originally identified Th1 with COVID severity. **c-d**, Violin plot and correlation analysis for scODIN-identified Th1 cells. **e-f**, Violin plot and correlation analysis for originally identified CD4-IL22 cells. **g-h**, Violin plot and correlation analysis for scODIN-identified Th22 cells. **i-k**, Violin plots of original CD4 naïve, CM, and EM cells comparing healthy controls to all COVID-19 patients. **l-p**, Violin plots of scODIN identified CD4 naïve, CM, and EM cells and double labels comparing healthy controls to all COVID-19 patients. **q-r**, Violin plots comparing healthy controls to all COVID-19 patients for q) original and r) scODIN identified total Tregs. **s-v**, Violin plots of scODIN identified Treg subsets comparing healthy controls to all COVID-19 patients. **w**, Violin plot comparing healthy controls to all COVID-19 patients for scODIN-identified CD4 CTL cells. Each dot corresponds to one patient or healthy control. Violin plot statistics: unpaired 2-tailed student’s t-test (rstatix). COVID-19 n = 80, healthy n = 21. Correlation analysis statistics: spearman’s rank correlation, smoothed conditional mean using method = “loess” and formula = “y ∼ x” with a 95% confidence interval. healthy n = 21, mild n = 23, moderate n = 30, severe n = 13, critical n = 14.

## Discussion

The identification of T cell subsets in scRNA-seq data has always been a difficult task. Biologically it is very simple to distinguish CD4 T cells from CD8 T cells, but due to few lineage-defining genes and many other overlapping genes, these cells are often indistinct on a UMAP. It has been even more difficult to distinguish smaller and rarer subsets within the CD4 T cell group due to dropout events of key identifying genes^7,8^. The most common approach in scRNA-seq analysis is clustering followed by naming the entire cluster based on a differential expression analysis compared to other clusters^15,23^. The caveat of this is that subsets may share many genes but a few key genes, such as FOXP3 or CXCR5, would play a dominant role in how they are annotated by an expert immunologist. For instance, we here clearly showed that what the authors of one study^32^ identified a large number of Th22 cells due to a very small fraction of cells in a cluster expressing IL22. While in that study the exact frequency of Th22 was not a major point, the improper identification of T cell subsets can lead to wrong conclusions, such as the Th22 example, being oppositely correlated after proper identification with scODIN. scODIN uses both a cluster-based approach on the top (PBMC) level, and a single-cell based approach on the detailed level such as CD4 T cells. Usually, computational methods use one or the other with scType^19^ and scCATCH^20^ using a cluster-based approach, and SCINA^21^ and scSorter^18^ using a single-cell based approach. Both approaches are valid, but for rare subsets the single-cell approach is more appropriate as they mostly do not form separate clusters. scType (cluster) and SCINA (single cell) performed equally well for two out of three datasets, while SCINA performed better on the third. Thus, it is more in the fine details of the approach, rather than the type of approach that matters. Some errors in subset identification came from initial over annotation, for example, scType labelling all CD8 cells as CD8 NKT and all B-cells as B-naïve. scODIN avoids this by simplifying the initial step to a single goal of correctly identifying main lineage populations rather than subsets. When dealing with large datasets like scRNA-seq, any operation that has to be calculated for every cell will slow down the process. scODIN and scType^19^ are almost equally fast at the calculation due to both using vectorized operations. However, scODIN still outperformed scType slightly, potentially due to a smaller and more streamlined database. The scType database has been generated in an unbiased manner from databases resulting in a large number of identifying genes, with every gene given the same weight, which is reduced if the gene is present in more than one subset. This differs from scODIN’s user-provided gene priority table, where it is possible to give a higher weight for very important genes, such as FOXP3 for Tregs or TBX21 (T-bet) for Th1 cells.

After initial clustering, the identification of rare cell subsets is a difficult task even for the expert annotator. In order to identify e.g. Tfr or CD4 CTL the data may need to be highly over-clustered before any cluster can be identified as these cell types. Over clustering complicates the annotation of the remainder of the subsets and makes the process very time-consuming. To overcome this, researchers have used manual “gating” on key markers to identify rare subsets, such as saying that Tregs should express FOXP3 and IL2RA or Th1 should express TBX21 and IFNG. This strategy is not unlike scODIN’s determination of core cell types, although writing manual code for each cell subset is time consuming and has been streamlined in scODIN. Additionally, scODIN enables expansion of the identified cells by virtue of kNN. Before kNN can be used, all core cell types must be annotated, as its principle is to search for nearest neighbors. If there are only a few cell types identified, the algorithm would generate false positive results, as there is little competition for being the nearest neighbor. On the other hand, if the reference dataset for the kNN is dominated by common cell types, they will also dominate the predicted cell identities. scODIN takes into account all of these things and expands the identified cell network in an as trustworthy way as possible while giving space for rare cells to be identified.

The example workflow in this report primarily focuses on CD4 T cell biology including Tregs, and we have thus fine-tuned the database for these cell types since this is our primary area of expertise. However, we also provide a basic template for all other immune subsets in PBMCs, using known markers from the literature. Due to its flexible design, the principle of scODIN is applicable to any cell type fine subsetting, and as new populations and markers are discovered, these can be simply added to database. In summary, scODIN enables a semi-holistic detailed cell type identification faster and more accurate than any other algorithm, mediating discovery of new biology and freeing up the hands of researchers to do more validation experiments.

## Methods

### scODIN gene priority table

The scODIN gene priority table has been generated from prior knowledge from the literature of canonical markers such as TBX21 (T-bet), CXCR5, and FOXP3 to separate CD4 T-cell subpopulations such as Th1, Tfh, and Tregs^10–14^. As cell types compete with each other for the cell identity, we found that fewer and more specific markers are more effective than a long list of less specific markers.

### scODIN score calculation

The scODIN score calculation is visually laid out in **Fig. S1**. As input scODIN needs a scaled gene expression table (in a Seurat object) and the gene priority table (**Fig. S1a**). Optional input is the tier column in the gene priority table and the accepted double labels table. Before the calculation begins, the gene names given in the gene priority table are checked for matching identities in the gene expression table. If there are gene names in the gene priority table that are not present in the gene expression table, the user is notified which gene names are missing. Next, to speed up the calculations, scODIN first subsets the gene expression table to genes in the gene priority table. For the calculation, the cell level is first defined, for example “Top” as top-level PBMC identification, or “CD4_T” if the identification should be done for CD4 T cells. Afterwards, a score for each cell type is calculated by taking the scaled expression value of each defining gene in the gene priority table and multiplying it with the user-defined gene priority. To adjust for the number of genes given in the priority table, the value is divided by the square root of the number of genes. Subsequently, one score for each cell type per cell is generated by taking the sum of each value for each gene. When the gene priority is positive, this will contribute to a higher scODIN score, but when the gene priority is negative it will contribute to a lower scODIN score. Next a user defined “core cell type cutoff” is applied to filter out low scoring cells, and a user defined “similarity threshold” to determine single- or double-labelled cells. The double-labelled cells will be tested against the “accepted double labels table”, and if accepted, the label is kept, and if not in the list, the label becomes “unknown” (**Fig. S1c**). If the number of labels above the “core cell type cutoff” is three or more, they are converted to “unknown” labels. In the kNN step, these unknown cell types are tested against the core cell types using a wider set of genes to better determine their dominating phenotype (**Fig. S1d**). Finally, the following information is added to the input Seurat object: single labels (containing only single labels), double labels (containing all double labels), and final labels (containing single and accepted double labels). Individual scODIN scores are also added to the column data for each cell type as the cell type name.

### scODIN tier system

The point of the scODIN tier system is to limit competition from common cell types allowing rare cells to be identified first. It breaks down the labelling of each cell type after the scODIN scoring has taken place, i.e. at the step where single labels and double labels are assigned. For the CD4 T cells in this report, we used 2 tiers: 1) Tfh, Tfr, nTreg, eTreg, T helper cells and CD4 CTL, and 2) naïve, CM and EM cells. Importantly, the top-level clustering-based identification was not tiered, but this could be implemented if desired.

### scODIN mixed and intermediate phenotype identifications

Identification of mixed and intermediate phenotypes is using the same code as the scODIN scoring system and accepted combinations are given in the “accepted double labels table”. In the current report, the table contains Th1/Th17, Th2/Th17, Tfh/Th1, Tfh/Th2, Tfh/Th17, eTreg/Th1, eTreg/Th2, eTreg/Th17naive/effector Treg, CD4-naive/CM, and CD4-CM/EM. The “accepted double labels table” may be adjusted as the user desires.

### kNN

The first part of kNN is to down-sample the reference core cells to a similar number using the “downsample_core()” function. This can be ignored, if the user wants to use all identified cells as the core reference. The kNN machine learning is conducted via a wrapper around Seurat’s FindTransferAnchors and TransferData^15^ (“odin_knn()”) with an additional function to implement user-defined score filters (“apply_score_filter()”). The user has the option to run an approximate nearest neighbor search by ANNOY (fastest) or an exact nearest neighbor search by RANN with eps (error bound) set to 0. Note, that changing eps from 0 would make RANN an approximate nearest neighbor search. Finally, the predictions are combined with the core cell types into one final Seurat object using “odin_merge()” function in scODIN to generate the “final_labels” column. It is furthermore possible to access where the cell label came from in the column data “source” as either knn or core.

### scODIN cluster-level cell type identification

Identifying many different cell types simultaneously is difficult, and one of the reasons behind scODIN’s tier system. On the top level, we would recommend identifying the major cell lineages using regular workflow clustering followed by scODIN top-level identification at the cluster level rather than the single cell level. To do so, the user may run “odin_cluster_scoring()” function in scODIN. The cluster labelling is based on a combined score considering the number of cells identified in a cluster of a specific cell type, as well as the scODIN score. The sum of all scODIN scores is multiplied by the square root of the number of cells. Results are sorted, and the highest-scoring cell type will be the cluster label.

### Comparison to other algorithms

The top PBMC-level identification of cell types has been previously reported by other groups with different algorithms^18–21^. To compare our results, we used these algorithms using their default settings. When a database was not available, we used our gene priority table formatted to fit the other algorithm’s requirements. For each algorithm, we cleared the RAM with gc() command, and recorded the start and end times, to calculate the run times. To estimate the percentage correctly assigned, we used the original author’s manual annotation as the ground truth. We considered a higher-level identification as correct, e.g. naïve CD4+ T cells are still CD4 T cells. However, we considered a lower-level identification as incorrect, e.g. all B cells cannot be classified as naïve B cells.

### Correlation analysis

Correlation analysis was done in R (v. 4.4.1) using cor.test from the R stats base packages on the 647k cell COVID-19 dataset^32^. Statistics were done by Spearman’s rank correlation, and smoothed conditional mean was plotted using method = “loess” and formula = “y ∼ x” with a 95% confidence interval.

### Pseudotime analysis

Pseudotime analysis was done in R using monocle3 (v. 1.3.7) using default settings and added to the Seurat object using AddMetaData. Results were plotted using VlnPlot grouped by the scODIN-determined cell identities.

### Version numbers

scODIN was made in R (v. 4.4.1) with dependencies Seurat (v. 5.1.0), readxl (v. 1.4.3), ggplot2 (v. 3.5.1), dplyr (v. 1.1.4), and scales (v. 1.3.0). Other algorithms used were scCATCH (v. 3.2.2), SCINA (v. 1.2.0), scSorter (v. 0.0.2), and scType (v. 1.0). For statistics, rstatix (v. 0.7.2) and base R statistics was used.

## Data availability

All data used in this work is publicly available. 3K dataset^3^: https://cf.10xgenomics.com/samples/cell/pbmc3k/pbmc3k_filtered_gene_bc_matrices.tar.gz, 161K dataset^15^: https://atlas.fredhutch.org/data/nygc/multimodal/pbmc_multimodal.h5seurat, 647K dataset^32^: https://covid19.cog.sanger.ac.uk/submissions/release1/haniffa21.processed.h5ad

## Code availability

The scODIN R package and bioinformatics data analysis pipeline have been deposited to GitHub (https://github.com/jonasns/scODIN). Full functionality requires the gene priority table available upon request.

## Acknowledgments

This work was supported by Japan Society for the Promotion of Science grant number 23K11304 (JNS), IFReC advanced postdoc program (DP), Japan Agency for Medical Research and Development, AMED grant number JP223fa627002 (JBW), Nippon Foundation FY2022 CiDER Cross-Departmental “Infectious Diseases” Research Promotion Program (JBW). This work was conducted as part of “The Nippon Foundation - Osaka University Project for Infectious Disease Prevention”.

## Author contributions

Methodology, J.T., D.P., and J.N.S.; Software, J.T., D.P., and J.N.S.; Validation, J.T., D.P., J.B.W., and J.N.S.; Formal Analysis, J.T., D.P., and J.N.S.; Investigation, J.T., D.P., and J.N.S.; Data Curation, J.T., D.P., J.B.W., and J.N.S.; Writing, J.T., D.P., J.B.W., and J.N.S.; Visualization, J.T., D.P., J.B.W., and J.N.S.; Conceptualization, J.N.S.; Supervision, J.B.W. and J.N.S.; Project Administration, J.B.W. and J.N.S.; Funding Acquisition, D.P., J.B.W., and J.N.S..

## Supplemental materials table of contents

**Figure S1:**
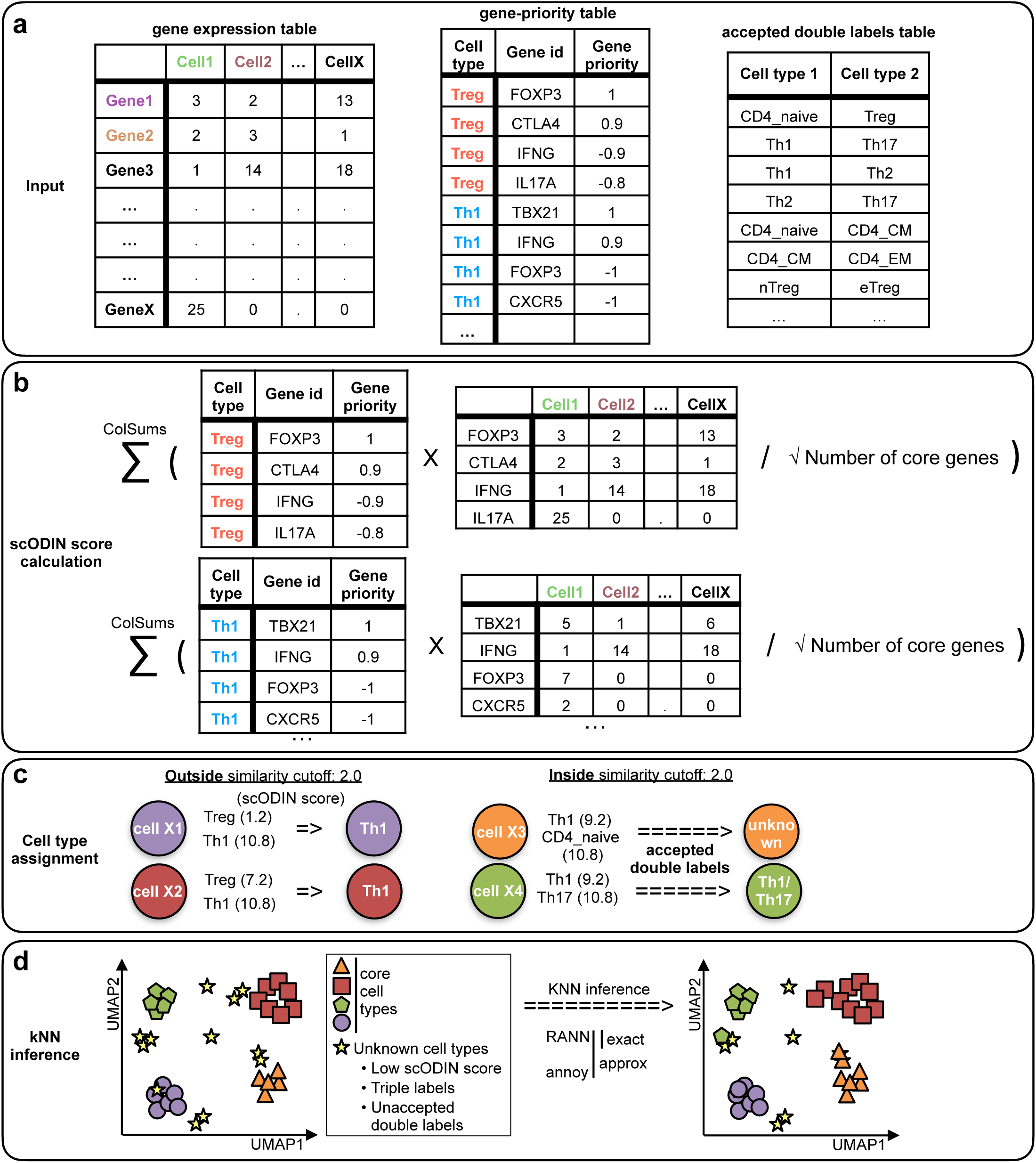
scODIN detailed overview. **a**, scODIN requires two inputs: a Seurat object with scaled gene expression values and the gene-priority table made in Excel (or similar). Optional input is an accepted double labels table. **b**, The scODIN score is calculated by taking the sum of a multiplication between the gene expression values with the gene priority table divided by the square root of the number of core genes. **c**, Examples of cell type assignment dependent on the similarity score and the accepted double labels. **d**, kNN inference is done using identified core cell types to predict the identity of unknown cell types. kNN can be done either by exact kNN (precise, slow) or approximate kNN (fast).

**Figure S2:**
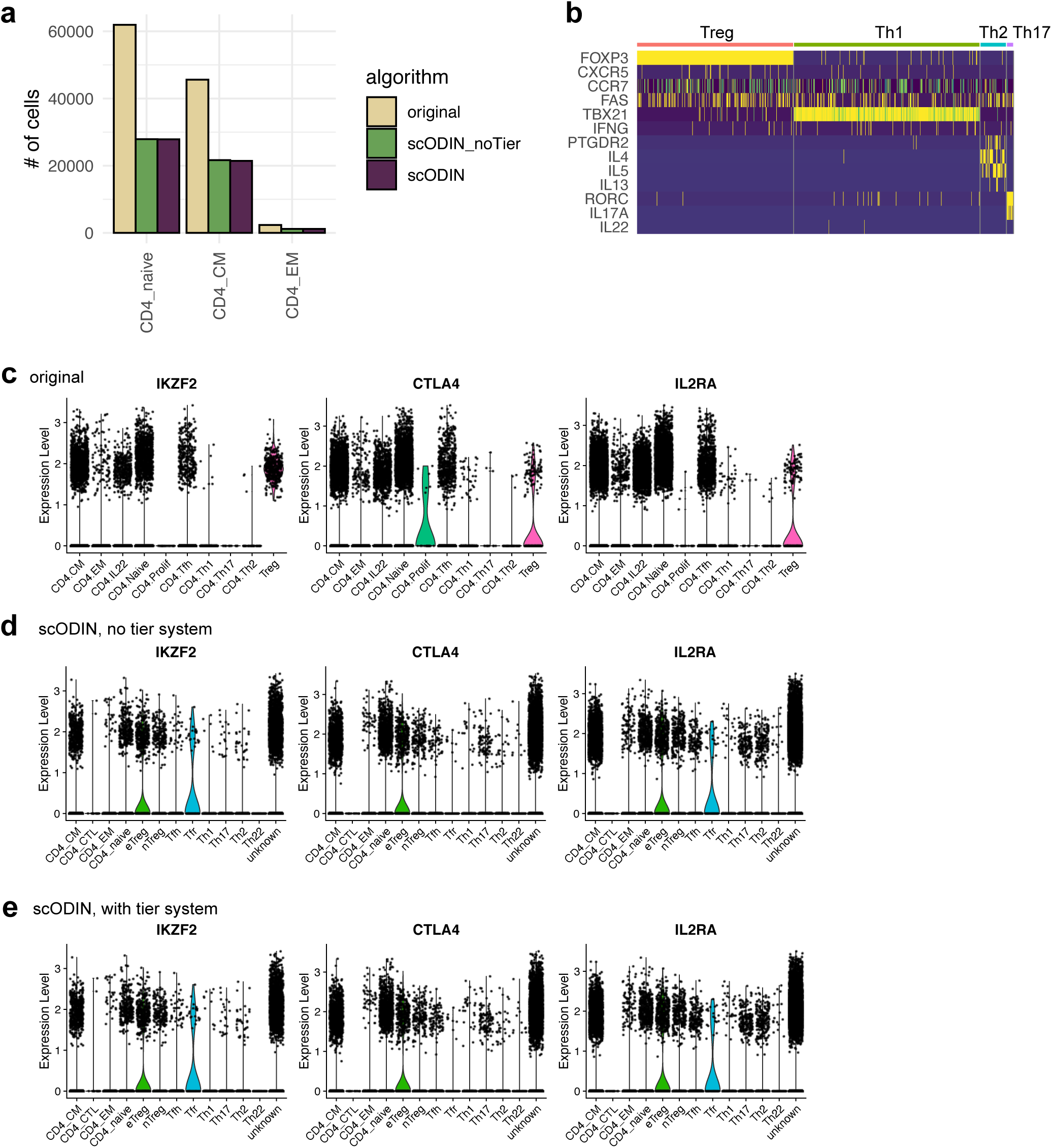
scODIN tier system additional figures. **a**, Number of generic cell types detected by scODIN +/- the tier system compared to the original annotation^32^. **b**, Heatmap of key markers for rare immune cell subsets for the original annotation (without dominating annotations: Tfh and Th22). **c-e**, Violin plots of IKZF2 (Helios), CTLA4, and IL2RA (CD25) expression for the original annotation (c), scODIN without tiers (d), and scODIN with tiers (e).

**Figure S3:**
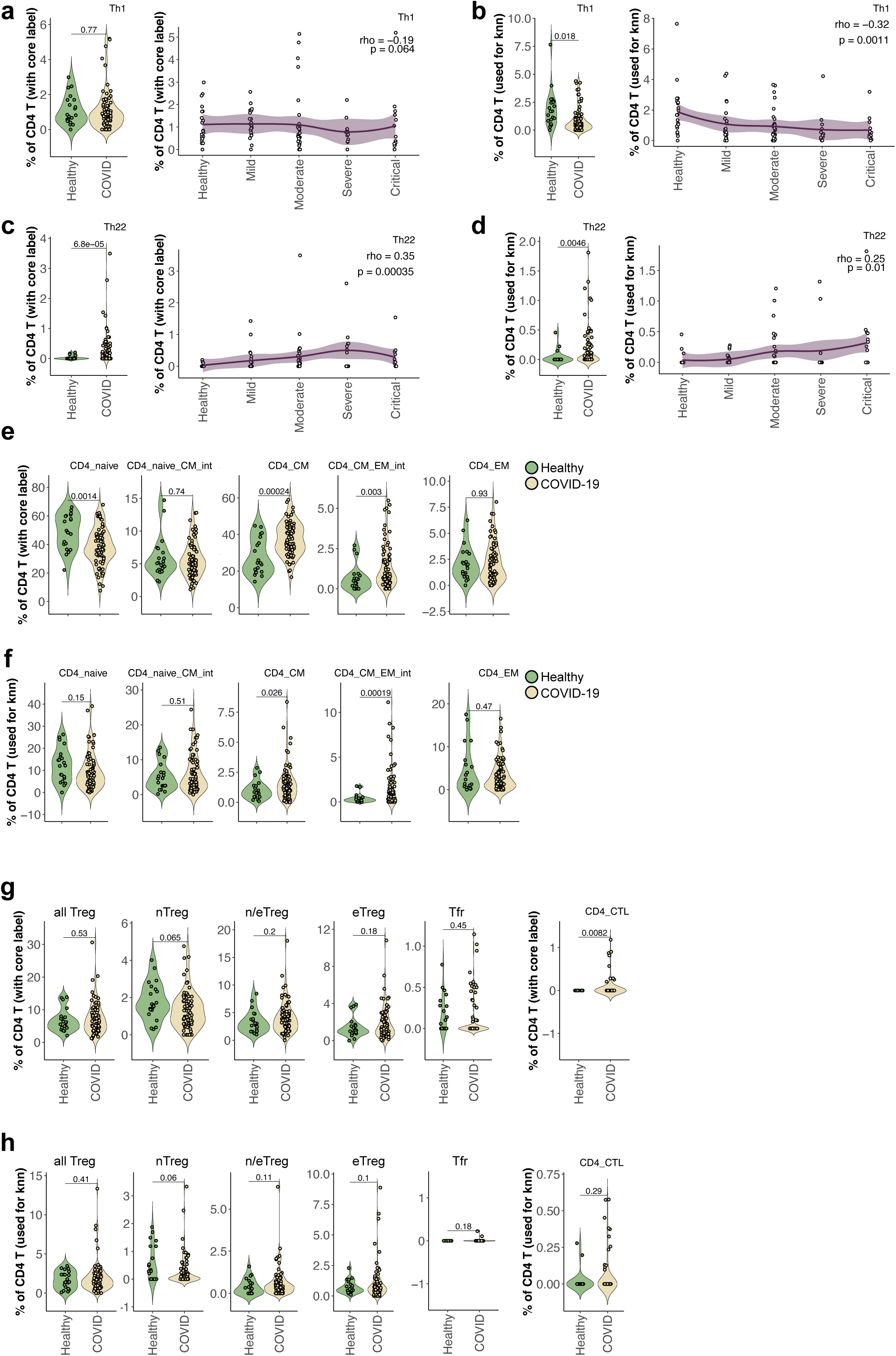
scODIN core and kNN annotations’ correlation with disease. **a**, Violin plot and correlation analysis comparing healthy controls to all COVID-19 patients for scODIN-identified core Th1 cells. **b**, Violin plot and correlation analysis comparing healthy controls to all COVID-19 patients for scODIN kNN-based identified Th1 cells. **c**, Violin plot and correlation analysis comparing healthy controls to all COVID-19 patients for scODIN-identified core Th22 cells. **d**, Violin plot and correlation analysis comparing healthy controls to all COVID-19 patients for scODIN kNN-based identified Th22 cells. **e**, Violin plots of scODIN identified core CD4 naïve, CM, and EM cells and double labels comparing healthy controls to all COVID-19 patients. **f**, Violin plots of scODIN kNN-based identified CD4 naïve, CM, and EM cells and double labels comparing healthy controls to all COVID-19 patients. **g**, Violin plots of scODIN identified core Treg subsets and CTL comparing healthy controls to all COVID-19 patients. **h**, Violin plots of scODIN kNN-based identified Treg subsets and CTL comparing healthy controls to all COVID-19 patients. Each dot corresponds to one patient or healthy control. Violin plot statistics: unpaired 2-tailed student’s t-test (rstatix). COVID-19 n = 80, healthy n = 21. Correlation analysis statistics: spearman’s rank correlation, smoothed conditional mean using method = “loess” and formula = “y ∼ x” with a 95% confidence interval. healthy n = 21, mild n = 23, moderate n = 30, severe n = 13, critical n = 14.

